# Altered nuclear envelope homeostasis is a key pathogenic event in C9ORF72-linked ALS/FTD

**DOI:** 10.1101/2024.02.01.578318

**Authors:** Riccardo Sirtori, Michelle Gregoire, Alicia Collins, Serena Santangelo, Bhavya Chatragadda, Robert Cullen, Antonia Ratti, Claudia Fallini

## Abstract

ALS and FTD are complex neurodegenerative disorders that primarily affects motor neurons in the brain and spinal cord, and cortical neurons in the frontal lobe. Although the pathogenesis of ALS/FTD is unclear, recent research spotlights nucleocytoplasmic transport impairment, DNA damage, and nuclear abnormalities as drivers of neuronal death. In this study, we show that loss of nuclear envelope (NE) integrity is a key pathology associated with nuclear pore complex (NPC) injury in *C9ORF72* mutant neurons. Importantly, we show that mechanical stresses generated by cytoskeletal forces on the NE can lead to NPC injury, loss of nuclear integrity, and accumulation of DNA damage. Importantly, we demonstrate that restoring NE tensional homeostasis, by disconnecting the nucleus from the cytoskeleton, can rescue NPC injury and reduce DNA damage in *C9ORF72* mutant cells. Together, our data suggest that modulation of NE homeostasis and repair may represent a novel and promising therapeutic target for ALS/FTD.

## INTRODUCTION

Amyotrophic lateral sclerosis and frontotemporal dementia (ALS/FTD) are fatal progressive neurodegenerative disorders affecting upper and lower motor neurons (MNs) in the brain and spinal cord, and cortical neurons in the frontal lobe in the central nervous system. Therapeutic treatments for ALS are extremely limited and do not significantly alter the course of the disease ^1^. In 90% of patients, death occurs within 3–5 years from clinical symptoms onset ^2^, highlighting the need for novel approaches and for a better understanding of the disease mechanisms. While the etiology of ALS is mostly unknown, genetic underpinnings can be identified in about 10% of cases, most commonly linked to mutations in the *C9ORF72, SOD1, TDP-43*, or *FUS* genes ^3^. The expansion of the six-nucleotide sequence GGGGCC in the first intron of the *C9ORF72* gene accounts for about 25-40% of familial cases and about 5% of sporadic ALS/FTD cases, making it one of the most common mutations throughout the whole patient population ^4^. The hexanucleotide repeat expansion (HRE) in *C9ORF72*, expanded to up to thousands of repeat units in ALS/FTD patients, forms nuclear and cytoplasmic RNA foci, and is translated into toxic dipeptides via repeat-associated non-AUG (RAN) translation ^5,6^. However, the mechanisms driving neuron death in this and other forms of ALS/FTD remain elusive.

Studies using *in vivo* and *in vitro* models of the disease have identified several cellular pathways associated with disease initiation and/or progression, including the accumulation of mislocalized and aggregating proteins such as TDP-43 ^7,8^, the activation of the stress and DNA damage response ^9,10^, injury to the nuclear pore complex (NPC), and nucleocytoplasmic transport (NCT) impairment ^11–13^. The NPC is a macromolecular protein assembly embedded in the double lipid bilayer of the nuclear membrane, and is the main gateway of macromolecular traffic between the nucleus and cytoplasm ^14,15^. Defects to NCT and NPCs are emerging as a central and driving mechanism of neurodegeneration not only in ALS/FTD, but in other neurodegenerative conditions such as Huntington’s and Alzheimer’s diseases ^12,16–20^. NPC injury alters RNA metabolism, protein localization, and cellular homeostasis, ultimately promoting cell death. While multiple pathways have been proposed to trigger and/or contribute to NPC injury in ALS/FTD, including oxidative stress and nucleoporin sequestration ^21,22^, nuclear envelope (NE) disrepair by the ESCRT-III complex ^23–25^, and cytoskeleton-induced nuclear damage ^20,26–28^, the underlying molecular mechanisms are still not clear.

Here, we investigated the role of mechanical stress on the nucleus as a significant driver of NPC and NE injury. Mechanical forces generated by the cytoskeleton can be transmitted onto the nucleus via the linker of nucleoskeleton and cytoskeleton (LINC) complex ^29–31^. This is a multiprotein complex embedded in the NE and is the main structure that links the nucleus to the plasma membrane. The LINC complex consists of the SUN1/2 proteins spanning the inner nuclear membrane, and Nesprin proteins embedded in the outer membrane ^29^. Cytoplasmic domains of nesprins bind cytoskeletal structures ^32^, while their KASH (Klarsicht, Anc-1, Syne homology) domains bind SUN proteins ^33^, which are in turn bound to the nuclear lamina and NPC through Emerin and NUP153 ^34–36^. While these interactions are required to maintain nuclear homeostasis in response to changes to the extracellular environment or intracellular state, they have been shown to also lead to transient NE ruptures ^36–38^, that are then repaired by the ESCRT-III complex ^39,40^. Interestingly, this activity was recently show to be facilitated by the ubiquitination and consequent removal of Nesprin2 from existing LINC complexes ^39^. This is accomplished by the ESCRT-III accessory protein BROX, a conserved Bro1 domain protein that is recruited to the sites of NE rupture by the ESCRT-III membrane repair machinery ^39^. BROX-dependent Nesprin2 removal locally relieves the NE from cytoskeleton-mediated pulling forces, facilitating the final closure of wounded nuclear membranes ^39^. In this study, we show that loss of NE integrity is a key pathology associated with NPC disruption in *C9ORF72* mutant neurons. We further show that relieving mechanical tension from the nucleus, either globally or locally around the NE rupture site, promotes NE repair and prevents the pathologic damage to the NPC trigger by the ESCRT-III complex in *C9ORF72* mutant cells, suggesting this could represent a novel therapeutic strategy for ALS/FTD.

## RESULTS

### NPC injury and NE breaks accumulate and co-occur in aged ALS/FTD neurons

Severe defects to NPCs have been extensively described in different models of ALS/FTD, including *in vivo* and *in vitro* models of *C9ORF72*-linked ALS ^26,41–44^. Terminally differentiated cells conversion into iPSCs erases almost completely the epigenetic memory and signs of aging of those cells ^45^, and neuronal differentiation driven by the ectopic expression of transcription factors (i.e., i^3^PSCs to i^3^Ns)^46,47^ accelerates significantly the definition of neuronal identity compared to methods based on sequential treatment with small molecules. Therefore, to observe the emergence of disease-relevant phenotypes in i^3^PSC-derived neurons, *in vitro* “aging” plays a fundamental role. To define the timing of NPC injury in cortical-like i^3^PSC-derived neurons (i^3^Ns) differentiated from mutant *C9ORF72* i^3^PSCs (C9^HRE^) compared to isogenic controls (C9^iso^) (**Figure S1**), we quantified the percentage of cells displaying altered NPC staining using two well-characterized NPC markers (i.e., mAb414 and RanGAP1) over the span of eight weeks (**Figure 1a-e**). We found that time played a key role in the emergence of NPC injury in C9^HRE^ i^3^Ns, which showed a gradual increase in the frequency of NPC disruption that became significant after 5 weeks *in vitro*, as judged from the qualitative analysis of the distribution of both markers at the NE (**Figure1 b, d**). This change was also accompanied by an overall reduction in mAb414 and RanGAP1 levels (**Figure 1 c, e; Figure S2**), indicating severe injury and loss of NPCs at the NE. Interestingly, we also found, using two independent i^3^PSC isogenic *C9ORF72* lines, that the spike in NPC disruption co-occurred with a significant increase in the frequency of ruptures or blebs of the NE, which led to the extrusion in the cytoplasm of nuclear DNA (**Figure 1 f-h**). Overall, these data suggested that NPC injury and loss of NE integrity are related and time-dependent events in mutant C9^HRE^ i^3^Ns.

**Figure 1.**
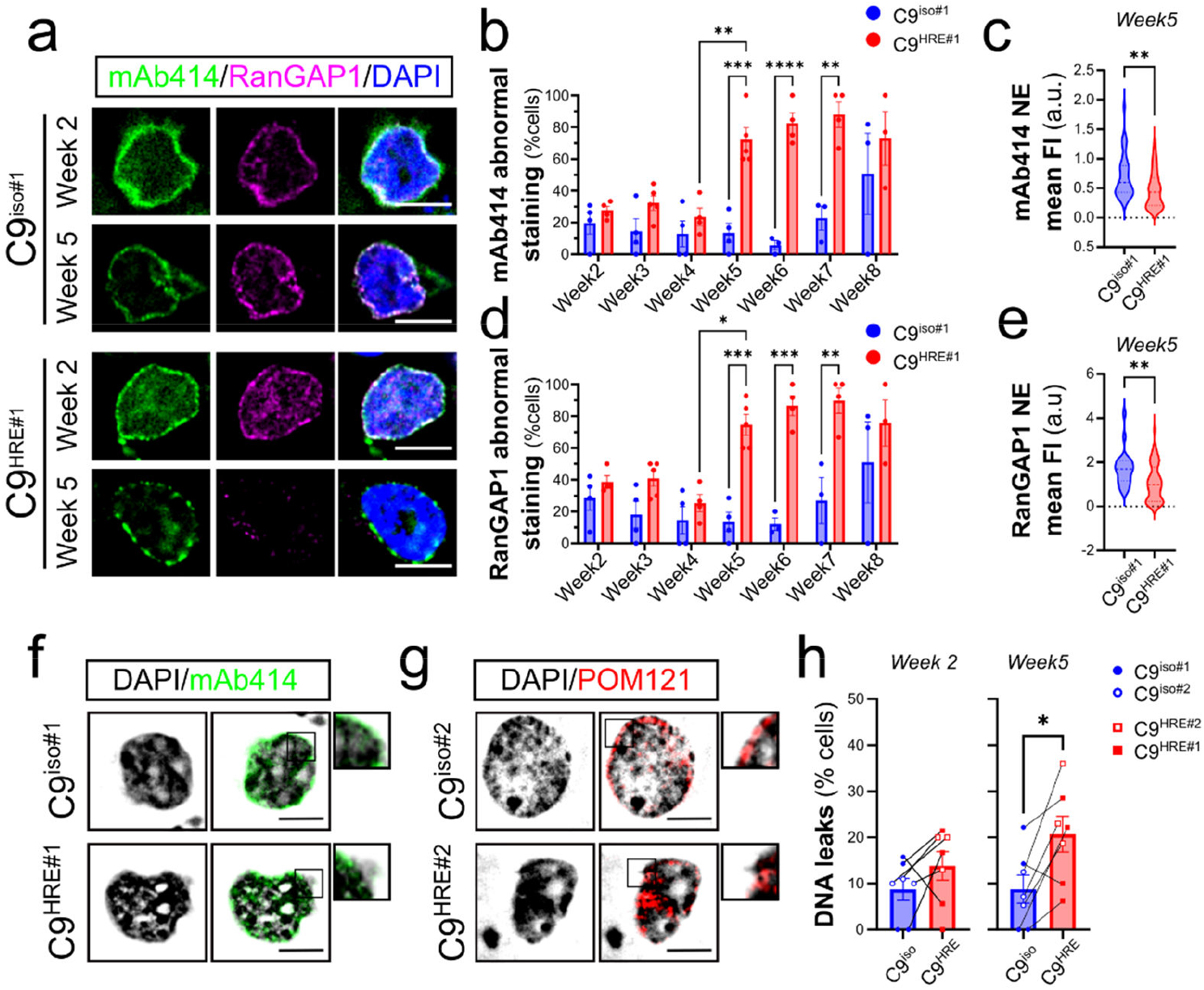
Age-dependent NPC disruption and NE rupture. **a**. Representative images of i^3^Ns derived from mutant (C9<SUP>HRE#1</SUP>) and isogenic (C9^iso#2^) iPSCs stained with the NPC markers mAb414 (*green*) and RanGAP1 (*magenta*) at 2 and 5 weeks *in vitro* after differentiation. DAPI (*blue*) labels the cell nucleus. **b-e**. Quantification of the percentage of cells with abnormal mAb414 (*b*) or RanGAP1 (*d*) staining was quantified every week between 2 and 8 weeks in culture. A significant increase in NPC disruption was evident around week 5 (two-way ANOVA, n=3-5 for all time points, ^***^ *p*<0.001). At the same time, the mean fluorescence intensity at the NE was significantly reduced in mutant cells compared to isogenic controls for both mAb414 (*c*) and RanGAP1 (*e*) (Welch’s t test, n=22 and 47 for C9^iso#1^ and C9^HRE#1^, respectively, ^**^ *p*<0.01). **f-h**. Five-week-old C9^HRE^ neurons display more frequent leaks of nuclear DNA in the cytoplasm (quantified in *h*) compared to isogenic controls (paired t test, n=7, ^*^ *p*<0.05). DAPI (*black & white* in *f* and *g*) labels the DNA, while mAb414 (*f, green*) and POM121 (*g, red*) were used to visualize the NE. For all, violin plots show data distribution with dashed lines representing mean and SD; bars represent mean and SEM. Scale bars: 10μm.

### The C9ORF72 hexanucleotide repeat expansion is sufficient to induce NCT defects and NE breaks in mitotic cells

While *in vitro* “aging” was required to induce NPC injury and NE ruptures in i^3^Ns, we wondered whether the presence of the *C9ORF72* hexanucleotide repeat expansion (HRE) alone was sufficient to cause similar defects in dividing cells, since their NE is subject to additional stresses during the cell division process. Thus, we transfected HEK293 cells with a plasmid expressing 80 repeats of the GGGGCC sequence [(G4C2)80] under the constitutive CMV promoter, as we previously described ^26^. Under these conditions, we found that the nucleocytoplasmic distribution of a GFP-NLS reporter, which should only localize to the nucleus due to the presence of a strong nuclear localization sequence (NLS), was severely altered in the presence of the (G4C2)80 repeat (**Figure 2a-b**). We also found that (G4C2)80-expressing cells were characterized by smaller and irregularly shaped nuclei (**Figure 2c-d**), suggesting alterations to the NE. In fact, an analysis of NPC and NE integrity showed a significant increase in the frequency of DNA leaks in the cytoplasm, accompanied by an overall reduction in the levels of the NPC-associated protein RanGAP1 (**Figure 2e-g**). Altogether, these results prove that the expression of the *C9ORF72* HRE is sufficient to disrupt nuclear morphology and nuclear envelope homeostasis in mitotic cells.

**Figure 2.**
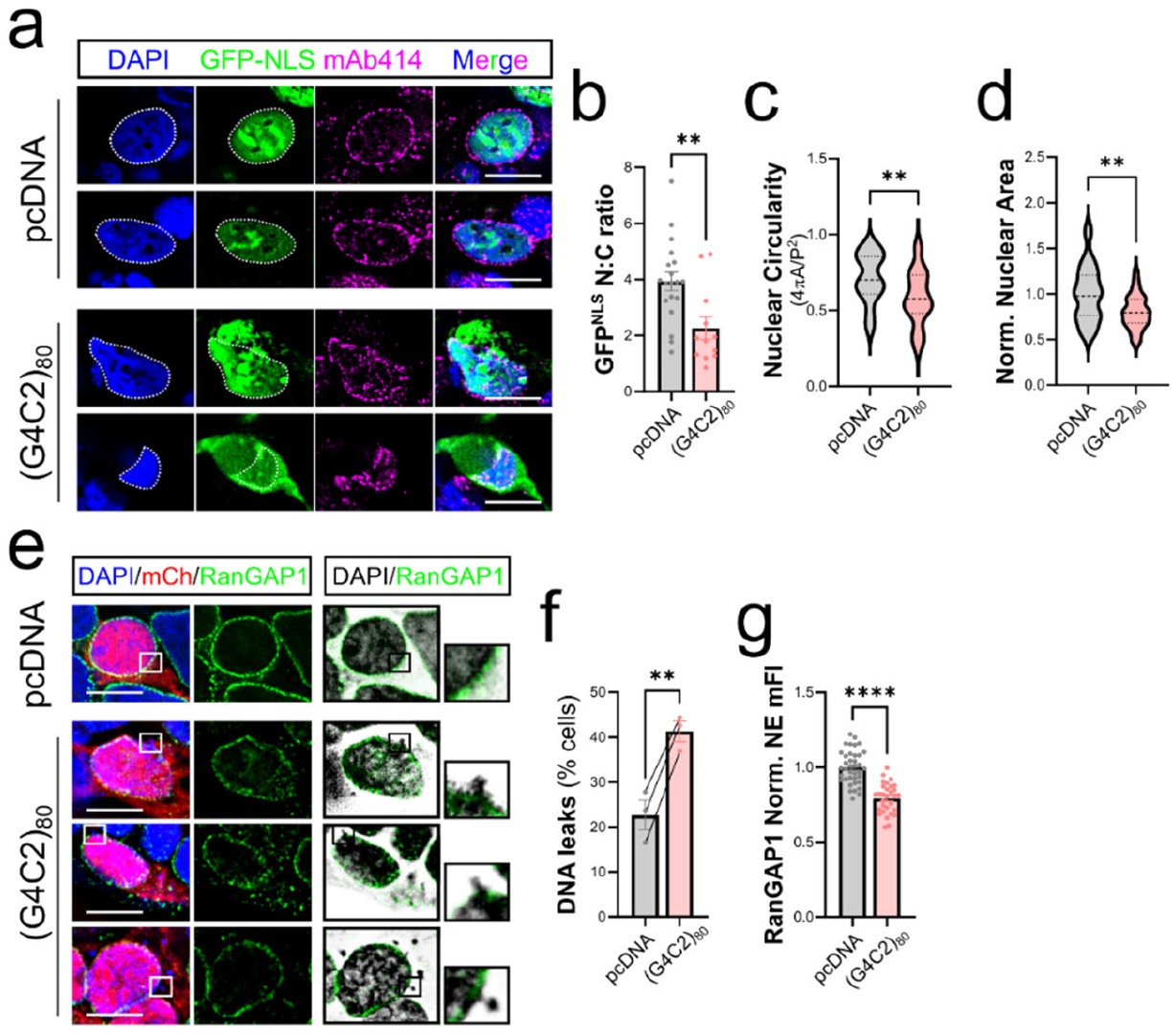
*C9ORF72* repeat expansion is sufficient to induce NPC injury and NE ruptures in HEK293 cells. HEK293 cells were transfected with the (G4C2)80 repeat expansion and compared to empty vector controls (pcDNA). The NPC markers mAb414 (*a, magenta*) and RanGAP1 (*b, green*) were used to visualize the NE. **a**.The nuclear fluorescent reported GFP-NLS (*green*) was co-expressed to monitor NCT efficiency. White dashed lines indicate the nuclear contour. **b**. The quantification of nucleus to cytoplasm (N:C) ratio of the GFP-NLS reporter shows a significant decrease in cells expressing the (G4C2)80 repeat, indicating cytoplasmic mislocalization. Mann Whitney tests; n=19 and 12 in pcDNA and (G4C2)80, respectively; ^**^*p*<0.01. **c-d**. The (G4C2)80 repeat also leads to less round and mishapen nuclei (*c*) with an overall smaller area (*d*). Student’s *t* test, n=35 and 31 for pcDNA and (G4C2)80, respectively, ^**^ *p*<0.01. **e-g**. Representative images show a significant increase in cytoplasmic DNA extrusions (quantified in *f*) accompanied by a reduction in RanGAP1 staining at the NE (quantified in *g*). Paired t test in *f*, n=3, ^**^p<0.01; Student’s *t* test in *g*, n=38 and 30 in pcDNA and (G4C2)80, respectively; ^****^*p*<0.0001. For all, violin plots show data distribution with dashed lines representing mean and SD; bars represent mean and SEM. Scale bars: 10μm.

### Cytoskeleton tension alters nuclear homeostasis leading to NE ruptures and DNA damage

Studies in several cellular systems have shown that NE damage can be triggered by alterations in the dynamic organization of the cytoskeleton via its interaction with the LINC complex ^36,48,49^. Of note, we have previously shown that positive or negative modulation of actin polymerization via pharmacological or genetic approaches impacts NPC integrity and affects nucleocytoplasmic distribution of shuttling proteins ^26^. We thus wondered if this pathway could be responsible for some of the phenotypes observed in the presence of the *C9ORF72* mutation. To answer this question, we first investigated the effects of increased actin polymerization on the nucleus of wild type HEK293 cells. Using a FRET-based assay ^50,51^ (**Figure S3a**), we quantified the tension exerted on the nucleus in HEK293 cells treated with IMM01, a known promoter of actin polymerization via the activation of formin function ^26,52^, or Latrunculin B, an actin depolymerizer. Interestingly, we found that while Latrunculin B caused an increase in FRET efficiency, indicative of reduced nuclear tension, IMM01 treatment led to a dose-dependent increase in nuclear tension (i.e., decrease in FRET efficiency) in fixed or live cells (**Figure 3a-b, S3b**). This increase in tension was also associated with a significant reduction in the levels of the nucleoporin POM121 at the NE (**Figure 3d**), an increase in DNA damage (i.e., increase in γH2AX levels, **Figure 3e**) and an increase in DNA leaks in the cytoplasm (**Figure 3f**), indicating an accumulation of NPC injury and loss of NE integrity. Interestingly, IMM01 treatment of 2-week-old C9^HRE^ i^3^Ns, a time point where no obvious NPC injury has yet occurred (*see Figure 1*), also led to the accumulation of DNA damage (**Figure 3g-h**), suggesting that this pathway is conserved in post-mitotic neurons and may be relevant to disease pathogenesis in ALS/FTD.

**Figure 3.**
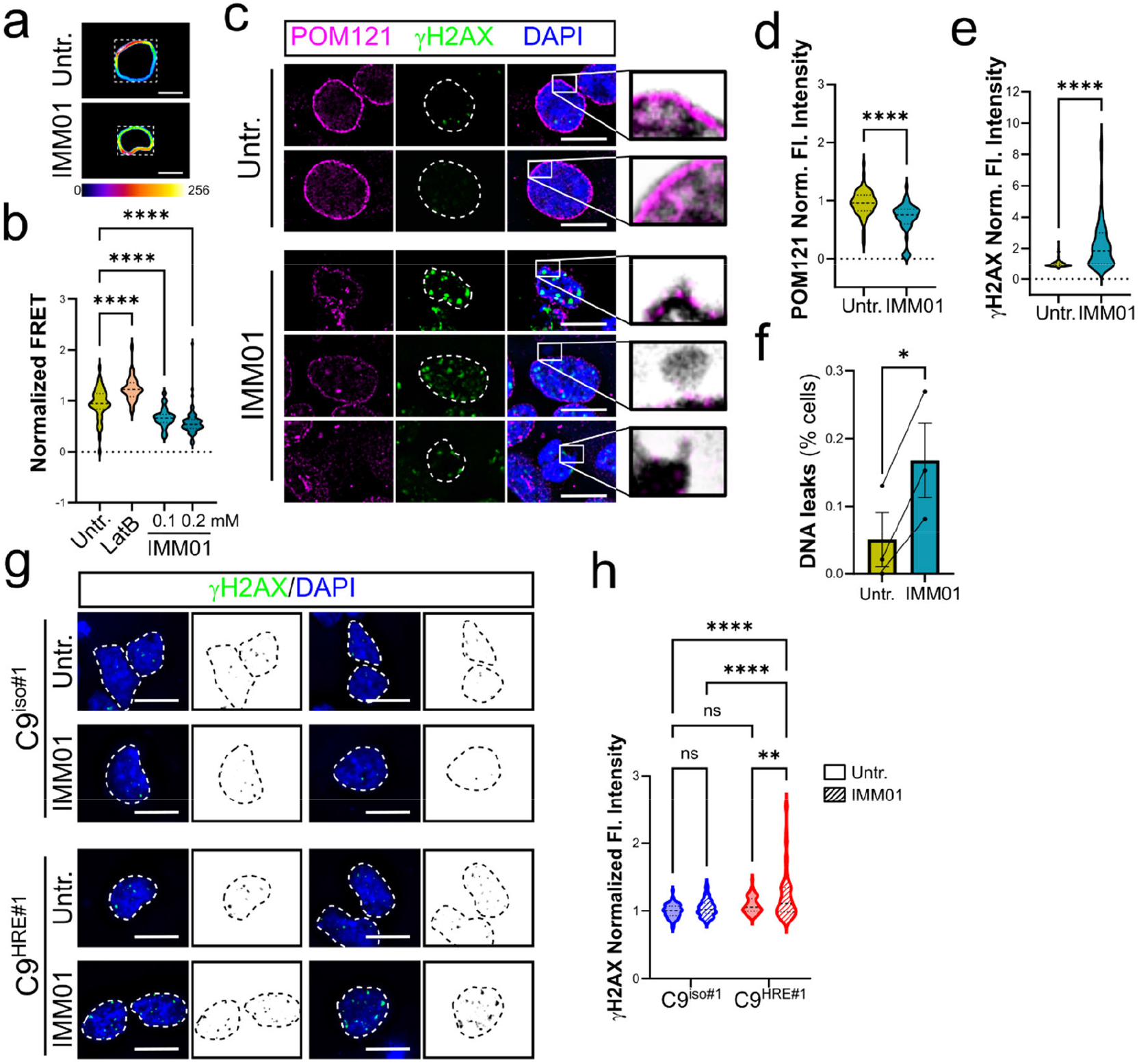
Nuclear injury is induced by IMM01 treatment in HEK293 cells and C9^HRE^ iNs. **a-b**. Representative images (*a*) and quantification (*b*) of nuclear tension exerted by the cell cytoskeleton using a FRET-based sensor in fixed HEK293 cells. Actin depolimerization using Latrunculin B (LatB, 1μM for 24 hours) caused an increase in FRET efficiency, indicating reduced tension, while IMM01 (0.1 or 0.2 mM for 5 hours), by promoting actin polymerization, led to a dose-dependent increase in nuclear tension, indicated by a decrease in FRET efficiency (one-way ANOVA, n=54, 34, 29, 53 for Untr, LatB, IMM01 0.1mM and 0.2mM, respectively; ^****^ *p*<0.0001). Dashed white squares in *a* indicate cropped area used for FRET data analysis. **c**. Representative images of HEK293 cells treated with 0.1mM IMM01 for 5 hours and stained with POM121 (*magenta*) to visualize NPCs, and γH2AX (*green*) to quantify DNA damage. DAPI (*blue*) was used to label DNA. **d-e**. POM121 levels at the NE are significantly reduced upon IMM01 treatment compared to untreated controls (*d*), while DNA damage is significanly increased (*e*) (Mann Whitney’s *t* test, n=55 and 47 for Untr. and IMM01, ^****^ *p*<0.0001. **f**. IMM01 treatement increases the percentage of cells with NE ruptures and cytoplasmic DNA leaks (paired *t* test, n=3, ^*^ *p*<0.05). **g-h**. IMM01 treatment (0.1mM for 5 hours) leads to increased DNA damage (γH2AX, *green* and *black&white* in *g*) in C9^HRE#1^ mutant i^3^Ns compared to isogenic controls (C9^iso#1^). DAPI (*blue*) was used to label nuclear DNA (two-way ANOVA, n= 69 (C9^iso#1^-Untr), 83 (C9^iso#1^-IMM01), 79 (C9^HRE31^-Untr), 80 (C9^HRE#1^-IMM01); ^**^ *p*<0.01,^****^ *p*<0.0001). Dashed lines in *c* and *g* define the nuclear contour. For all, violin plots show data distribution with dashed lines representing mean and SD; bars represent mean and SEM. Scale bars: 10μm.

To further demonstrate that excessive actin-based forces exerted on the NE are directly linked to the observed alterations of nuclear homeostasis, NPCs injury, and DNA damage, we employed a complementary strategy that relied on the use of a dominant negative form of the LINC protein Nesprin2 (KASH^DN^) ^32^. This well-established chimeric construct lacks the ability to bind to actin filaments because its actin-binding domain has been replaced by the red fluorescent protein mCherry. However, it can efficiently incorporate into existing LINC complexes at the NE via the essential SUN-binding domain. These interactions effectively cause the cell’s nucleus to become disconnected from the actin cytoskeleton ^32,53,54^. We found that expression of the KASH^DN^ in HEK293 cells prevented the accumulation of nuclear damage following IMM01 treatment compared to cells transfected with mCherry alone, as shown by a restoration of nuclear size, nucleocytoplasmic distribution of the essential shuttling factor RAN ^55^, and a significant reduction in the frequency of NPC injury (**Figure 4a-d**), proving that NE tension plays an important role in cytoskeleton-mediated nuclear damage. Given these promising results, we used the same strategy to attempt to rescue *C9ORF72* HRE-induced nuclear damage. Co-transfecting KASH^DN^ and the (G4C2)80 constructs resulted in a significant normalization of nuclear shape and reduced the frequency of (G4C2)80-expressing cells with severe NPCs alterations (**Figure 4e-g**). Overall, these results strongly support the hypothesis that mechanical strain on the nucleus significantly contributes to NPC pathology in *C9ORF72* mutant cells.

**Figure 4.**
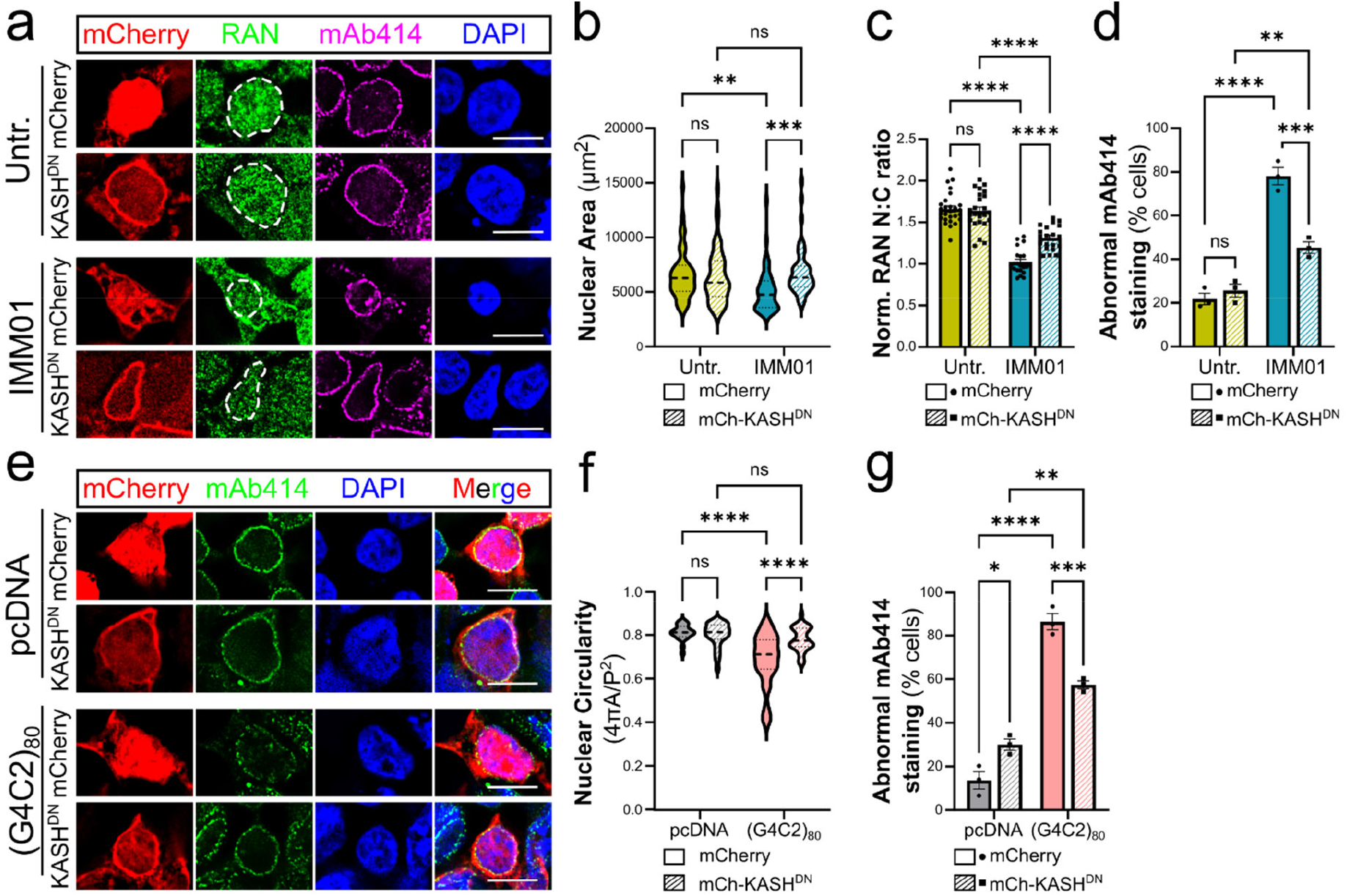
Actin-induced mechanical strain on the nucleus mediates nuclear abnormalities and NPC injury via the LINC complex. **a**. Representative images of HEK293 cells transfected with the KASH<SUP>DN </SUP>construct or mCherry as control (*red*) and treated with 100μM IMM01 for 5 hours. Cells were stained with antibodies to the nuclear shuttling transport factor RAN (*green*) and the NPC marker mAb414 (*magenta*). **b-d**. Quantification of nuclear size (*b*), RAN nucleus to cytoplasm (N:C) ratio (*c*), and frequency of NPC injury (*d*) in transfected cells with or without IMM01 treatment. IMM01 treatment significanly altered all three parameters in mCherry-transfected cells, but its effects were blunted by the presence of the KASH^DN^ contruct (two-way ANOVA, n=50 for all in *b*; n=27, 24, 21, and 20 in order of graph appearance in *c*, and n=3 each in *d*; ^**^ *p*<0.01,^***^ *p*<0.001,^****^ *p*<0.0001). **e**. HEK293 cells were co-transfected with the (G4C2)80 and KASH^DN^ (*red*) contructs. An empty vector (pcDNA) and mCherry alone functioned as negative controls. The NPC was visulaized using the mAb414 marker (*green*). **f-g**. Quantification of nuclear circularity (*f*) and NPC injury (*g*) shows a significant rescue of the damage induced by the (G4C2)80 contruct in the presence of the KASH^DN^ chimeric protein. (two-way ANOVA, n=20, 26, 27, and 35 in order of graph appearance in *f*, n=3 each in *g*; ^**^ *p*<0.01,^***^ *p*<0.001,^****^ *p*<0.0001). DAPI (*blue*) was used to visualize the cell’s nucleus in *a* and *e*. The white dashed lines in *a* highlight the nuclear contour. For all, violin plots show data distribution with dashed lines representing mean and SD; bars represent mean and SEM. Scale bars: 10μm

### The ESCRT-III associated factor BROX can restore NPC integrity and normalize ESCRT-III nuclear accumulation

Under normal conditions, NE ruptures induced by mechanical strain and/or normal cellular processes are remodeled and repaired by ESCRT-III machinery with the recruitment of the AAA ATPase VPS4 ^40,56,57^. Recent reports have shown that in *C9ORF72* mutant neurons, the ESCRT-III complex components CHMP7 and VPS4 pathologically accumulate in the nucleus, driving excessive membrane remodeling and NPC loss ^23–25^. Given our data showing that uncoupling cytoskeletal tension from the nucleus could revert *C9ORF72*-induced nuclear alterations, we wondered whether the BROX-dependent reduction of NE tension focally at the site of rupture could prevent the toxic accumulation of ESCRT-III complex components and subsequent injury to NPCs.

Indeed, we found that siRNA-mediated downregulation of endogenous BROX led to enhanced NPC injury in both control and mutant cells (**Figure 5a-c**), proving the essential homeostatic role of this process in nuclear maintenance. Importantly, overexpression of GFP-tagged BROX led to a significant reduction in NPC injury, and to the restoration of the nucleocytoplasmic distribution of RAN in (G4C2)80-expressing cells (**Figure 5d-f**), without causing any negative outcomes in control cells. These positive effects depended on BROX-Nesprin2 interaction, as the introduction of the L350A mutation in BROX, which renders it unable to bind Nesprin2 ^39^, completely abrogated the rescue (**Figure S4**). Finally, we investigated whether the protective effects of BROX overexpression on the accumulation of NPC injury depended on the modulation of VPS4 and CHMP7 nuclear recruitment. To that end, we fractionated the nuclear and cytoplasmic proteomes from (G4C2)80 and control cells with or without GFP-BROX overexpression and quantified VPS4 and CHMP7 relative abundance in both fractions by western blot. As expected, we observed the accumulation of VPS4 in the nuclear fraction of (G4C2)80 cells compared to controls, which was significantly reduced by GFP-BROX overexpression (**Figure 5g-i**). A similar trend was observed for CHMP7, albeit it did not reach statistical significance. Altogether, these results highlight the role of the LINC-actin interaction in driving NPC injury in mutant *C9ORF72* cells and prove that fostering NE repair through the reduction of the NE tension effectively rescues NPC injury induced by *C9ORF72* HRE overexpression, possibly by promoting the clearance of VPS4 and CHMP7 from the nucleus.

**Figure 5.**
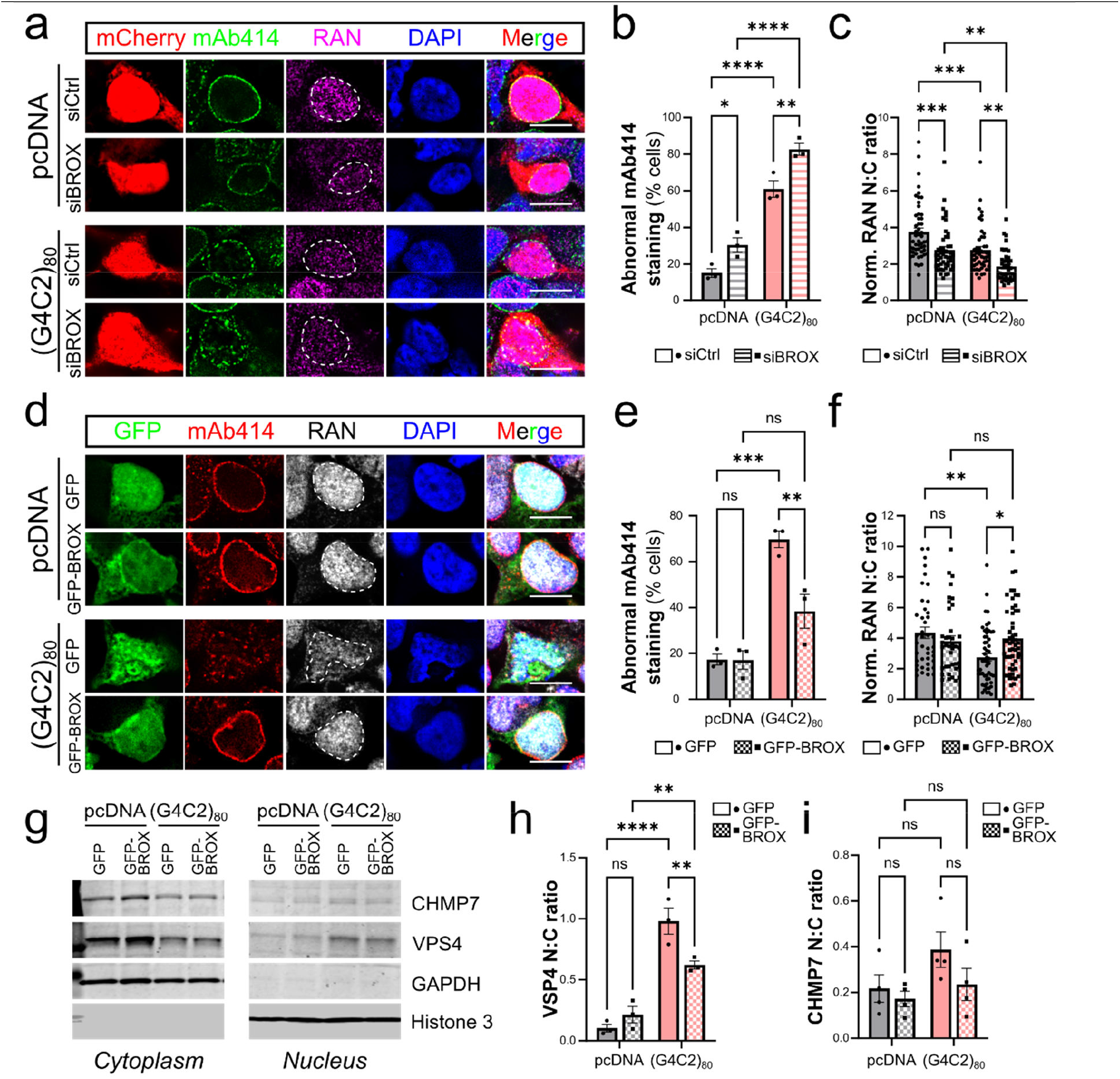
BROX cellular levels modulate NPC injury via the ESCRT-III complex. **a-c**. Representative images (*a*) and quantification of NPC injury (*b*) and RAN N:C distribution (*c*) in control or (G4C2)80 HEK293 cells transfected with siRNA targeting BROX (siBROX) or with a non-targeting control (siCtrl). Knock down of endogenous BROX was sufficient to induce NPC injury (mAb414, *green*) and RAN mislocalization (*magenta*) in control cells and exacerbated the toxicity of the *C9ORF72* HRE in (G4C2)80 cells (two-way ANOVA; n= 3 for each in *b*; n=52, 47, 47, 43 in order of graph appearance in *c*; ^*^*p*<0.05, ^**^*p*<0.01, ^***^*p*<0.001, ^****^*p*<0.0001). **d-f**. Representative images (*d*) and quantification of NPC injury (*e*) and RAN N:C distribution (*f*) in control or (G4C2)80 HEK293 cells transfected with GFP-BROX or GFP alone (*green* in *d*). Overexpression of GFP-BROX significantly rescued NPC injury (mAb414, *red*) and RAN mislocalization (*grays*) induced by the presence of C9ORF72 (G4C2)80 repeat expansion (two-way ANOVA; n=3 in *e*; n=42, 39,55, 52 in order of graph appearance in *f*; ^*^*p*<0.05, ^**^*p*<0.01, ^***^*p*<0.001; *n*.*s*., not significant). DAPI (*blue*) was used to visualize the cell’s nucleus in *a* and *d*. The white dashed lines in *a* and *d* highlight the nuclear contour. Scale bars: 10μm. **g-i**. Representative blot (*g*) and quantification of VPS4 (*h*) and CHMP7 (*i*) levels in the cytoplasmic (*left*)and nuclear fraction (*right*) of control (i.e. pcDNA) or (G4C2)80 HEK293 cells transfected with GFP-BROX or GFP alone. GAPDH and Histone 3 were used as fraction-specific loading controls (two-way ANOVA; n=3 in h, n=4 in I; ^**^*p*<0.01, ^****^*p*<0.0001; *n*.*s*., not significant). For all, bars represent mean and SEM.

### Fostering NE repair, BROX expression rescues HRE-induced DNA damage

Since we have previously shown that NPC injury associates with cytoplasmic DNA leaks and increased DNA damage in mutant *C9ORF72* cells, we wondered if promoting NE repair via GFP-BROX overexpression would be sufficient to prevent such toxic phenotypes. Thus, we co-expressed GFP-BROX in (G4C2)80 or control cells (i.e., pcDNA) and quantified the frequency of DNA extrusions in the cytoplasm by immunofluorescence and the accumulation of DNA damage by western blot analysis of the levels of the γH2AX marker (**Figure 6**). As expected, we found that the presence of the G4C2 repeat expansion led to an increase in the percentage of cells displaying DNA leaks, which was associated with higher levels of γH2AX, indicative of increased DNA damage. However, the expression of GFP-BROX returned both parameters within the levels of control cells. Overall, these data suggest that changes in nuclear tension can significantly contribute to disease-relevant alterations, and further support the importance of NE repair mechanisms in modulating nuclear homeostasis and function.

**Figure 6.**
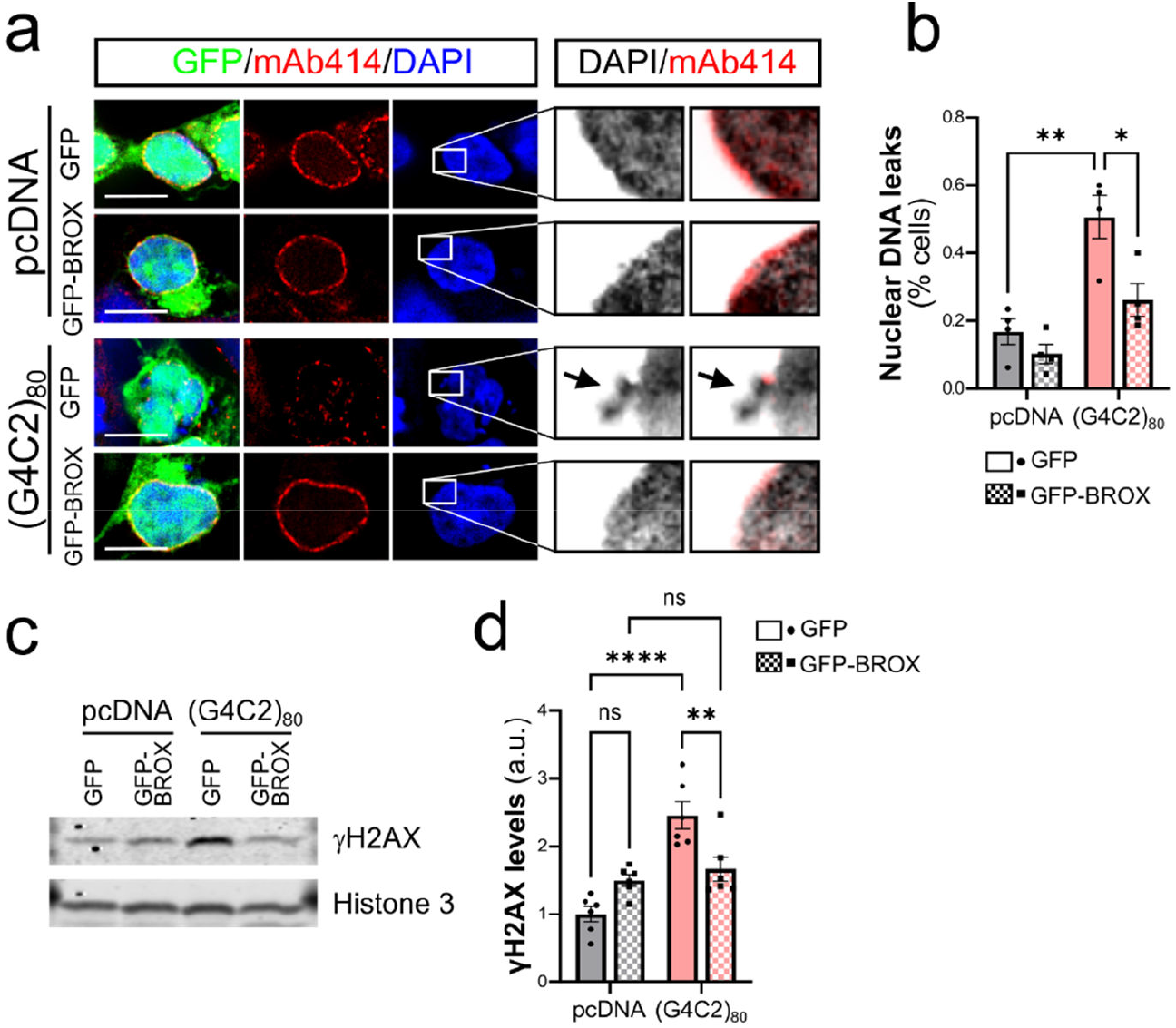
BROX promotes NE repair and reduces DNA damage in *C9ORF72* mutant cells. **a**. Representative images of HEK293 cells transfected with the empty vector pcDNA or the *C9ORF72* HRE (G4C2)80 and co-expressing either GFP or GFP-BROX (*green*). The NPC marker mAb414 (red) was used to visualize the NE. DAPI (*blue* in left panel, *black&white* in right panel) labels nuclear DNA. **b**. Quantification of the percentage of cells with DNA leaks shows a significant increase in (G4C2)80-expressing cells that is returned to control levels with the co-expression of GFP-BROX. **c-d**. Representative western blot and quantitative analysis of γH2AX levels in control pcDNA or (G4C2)80 cells co-expressing GFP or GFP-BROX. Histone 3 was used as loading control. Bars represent mean and SEM; two-way ANOVA; n=4 in *b*, n=5 in *d*; ^*^*p*<0.05, ^**^ *p*<0.01, ^****^ *p*<0.0001; n.s.= non-significant. Scale bars: 10μm.

## DISCUSSION

ALS is a complex disease with unknown etiology, and mutations in a wide array of genes have been associated with its pathogenesis ^58–60^. Although a common mechanism or comprehensive model linking these different gene mutations and sporadic disease features has not yet emerged, mounting evidence points to the dysregulation of nuclear homeostasis as a prime candidate. First, injury to NPCs has been described as a key pathologic event in numerous *in vitro* and *in vivo* models of ALS, importantly including patients’ postmortem tissue ^11,12^. Second, these defects have been associated with structural and morphological changes to the NE and nuclear lamina in most, although not all, models ^19,20,26,61^, reminiscing of some of the defects observed in laminopathies such as progeria syndrome. Changes to NE integrity in the context of neurodegenerative diseases can be considered significant triggers of disease progression, as they may initiate a cascade of events, including ESCRT-III recruitment and NPC injury ^23,24^, loss of segregation of nuclear and cytoplasmic content, and consequent accumulation of DNA damage ^62^. Loss of NE integrity has also been proposed to induce or promote protein aggregation, as recently shown in a tau model of FTD ^61^. Third, changes in nuclear and nucleolar size, a tightly regulated feature that depends on the complex interaction between the nuclear lamina, LINC complex, cytoskeleton, and chromatin ^38,49^, have been described in human postmortem tissues ^63^, along with broad alterations to chromatin organization and epigenetic regulation ^64,65^. Overall, these observations suggest that changes to the general homeostasis of nuclear structure and function may be key events in ALS pathogenesis, although the mechanisms and molecular players involved remained undefined.

In this study, we explored how nuclear dis-homeostasis is mechanistically linked to changes to mechanical forces exerted onto the nucleus by the cell’s cytoskeleton. Using mutant *C9ORF72* iPSC-derived neurons and (*G4C2*)80-overexpressing HEK293 cells, we showed that these different cell types and ALS models share the pathologic feature of nuclear defects, which depends on both the genetic background of the cells and the *in vitro* aging process. The defects we comprehensively describe here go beyond the previously characterized alterations in NPC integrity, involving striking changes to the whole nuclear size and shape, and to the capacity of the NE to compartmentalize the nucleus, leading to protein mislocalization, DNA leaks into the cytoplasm, and increased DNA damage.

Interestingly, many of the processes we found altered in the presence of the *C9ORF72* mutation are known to be mediated by the association between the NE and the cytoskeleton via the LINC complex. The tensile forces exerted on the nucleus by the actin cytoskeleton are in fact key to stabilizing and maintaining the nuclear morphology, as they compensate inward pulling forces generated by the heterochromatin bound to the nuclear lamina ^38,66,67^. An imbalance between these forces can lead to the deformation of the nucleus, which can then promote NE rupture ^67^. Our previous work investigating the pathogenic mechanisms leading to neurodegeneration in the presence of mutant PFN1 had identified the actin cytoskeleton as a modulator of NPC integrity ^26^. Specifically, we found that positive or negative disturbances to actin homeostasis were associated with NPC disruption and nuclear import deficits. Here, we expanded those observations to show that cytoskeletal tension is directly and mechanistically connected to NE alterations and NPC injury. In fact, we found that enhanced actin polymerization mediated by IMM01, a potent formin agonist ^26,52^, led to a significant increase in the mechanical tension at the NE, which triggered NE weakening and rupture, NPC injury, cytoplasmic DNA leaks, and increased DNA damage. Furthermore, we found that disconnecting the NE from the cytoskeleton using a dominant-negative form of the LINC complex component Nesprin2 (i.e. KASH^DN^) ^32,53^ was sufficient to rescue such alterations. In fact, we found that KASH^DN^ expression could prevent changes to nuclear size and shape, as well as NPC disruption caused by cytoskeletal forces induced by both IMM01 treatment and the expression of the *C9ORF72* HRE.

NE breaks that occur as a consequence of normal or pathological cellular process are repaired by the endosomal sorting complex required for transport (ESCRT)-III complex. The chromatin-associated protein barrier to autointegration factor (BANF1) and inner nuclear membrane proteins such as LEMD2, which become exposed to the cytoplasm because of a NE rupture, recruit CHMP7 to induce the local polymerization of ESCRT-III filaments, leading to NE resealing through the action of the AAA-ATPase VPS4 ^62,68,69^. A recent study from Dr. Agromayor’s group further showed that relaxation of nuclear envelope tension at the rupture point is a fundamental step for the resealing of NE ruptures ^39^. This reduction in NE tension is achieved thanks to the action of the ESCRT-III associated factor BROX, which promotes Nesprin2 ubiquitination and consequent degradation at the rupture point, thereby reducing the transmission of cytoskeletal forces that may prevent the efficient NE repair ^39^. Interestingly, aberrant nuclear accumulation of the ESCRT-III components CHMP7 and VPS4 has recently been identified as an upstream pathophysiological event that can trigger NPC injury and contribute to downstream TDP-43 dysfunction in sALS and C9-ALS ^23–25,70^. In this study, we show that, similar to what we observed following IMM01 treatment, BROX siRNA-mediated downregulation exacerbated nuclear alterations caused by the overexpression of the *C9ORF72* HRE. More importantly, we found that the overexpression of GFP-tagged BROX fostered the efficient repair of NE integrity, promoting the clearance of VPS4 and CHMP7 from the nucleus and thereby preventing NPC injury. While downregulation of CHMP7 and VPS4 has been proposed as a potential novel therapeutic avenue for ALS/FTD, the ESCRT-III complex remains a fundamental machinery required for the maintenance of NE integrity, among other equally important cellular functions ^57^. Our study shows that reducing the ESCRT-III complex dwell time on the NE by increasing NE repair efficiency may be a better strategy to prevent NPC injury, without affecting the general activity of the ESCRT-III machinery.

In conclusion, our study clearly demonstrates that NE alterations are a key pathologic feature of ALS/FTD using different cellular models. We found that increased mechanical stress exerted on the nucleus significantly contributes to NPC injury and NE weakness, and that releasing the nucleus from such cytoskeleton-dependent tension, either globally (i.e. KASH^DN^) or focally (i.e. GFP-BROX), can prevent NE disruption. While our data suggest a direct connection between HRE expression, mechanical stress on the nucleus, and NE alterations, whether and how mutations in the *C9ORF72* locus alter cytoskeletal homeostasis remains to be elucidated. It is in fact possible that the accumulation of NE injury may occur downstream of *C9ORF72*-induced DNA damage, as it has been shown in the context of cancer ^71^. In that case, the protective effect of KASH^DN^ and GFP-BROX expression would act through a parallel and complementary pathway. While further studies will be necessary to fully address these questions, our current results nonetheless suggest that NE ruptures trigger NPC injury and NCT disruption in *C9ORF72* mutant cells, and that these phenotypes can be modulated by relieving cytoskeleton-induced strain from the nucleus, opening new avenues for therapeutic development.

## STAR Methods

### i^3^PSC culture and differentiation

i^3^PSCs were grown in StemFlex media and maintained with daily media changes on Matrigel-coated plastic dishes. Colonies were passaged using 0.5mM EDTA in Ca^2+^- and Mg^2+^-free 1x PBS every 4-6 days, and media was refreshed daily. ROCK inhibitor (10μM; Y-27632, SellChem) was added after each passage for 24 hours.

To generate i^3^PSCs, we used the method described by Dr. Ward with minimal changes ^46,47^. This method relies on the integration of a gene expression cassette in the safe-harbor CLYBL1 locus and produces >90% pure neuronal populations. For this study, we used two pairs of isogenic lines (**Table S1**), in which the differentiation cassette containing the NGN2 transcription factor, mApple^NLS^ fluorescent protein, and blasticidin resistance gene (Addgene plasmid # 124229) were inserted using CRISPR/Cas9 into the CLYBL safe harbor locus as described ^46,47^. Guide RNAs for CRISPR/Cas9 integration of the cassette were obtained from Synthego and designed using their dedicated software. iPSCs were transfected with the plasmid containing the differentiation cassette and ribonucleoprotein particles containing the nuclease Cas9 and the specific gRNAs using Lipofectamine ES according to the manufacturer’s recommendations. Selection of positive clones was accomplished using blasticidin (10μg/ml) treatment. In addition, mApple^NLS^-positive clones were visually selected to increase the purity of the population. To minimize the effects of inter-clonal variation, all cells positive for the cassette integration were pooled together and expanded. To induce differentiation of i^3^Ns, 2 μg/ml doxycycline (Millipore Sigma) was added to the medium for 2 days, while proliferating cells were killed off with 40μM BrdU treatment. Cells were than plated on poly-ornithine and poly-lysine (1mg/ml; Millipore Sigma) coated coverslips and switched to neuronal medium (Neurobasal, 1% GlutaMax, 2% B27, 1% N2, 1% NEAA, 1μg/ml laminin). To verify full differentiation, positivity for Map2 and Tau and negativity for Oct4 and Sox2 was assessed. All reagents were acquired from Thermo Fisher Scientific unless otherwise stated.

### HEK293 cell culture and transfection

HEK293 cells were grown in DMEM media supplemented with 10% FBS. For immunofluorescence experiments, cells were plated on poly-lysine coated coverslips (1mg/ml; Millipore Sigma) and allowed to recover for 24 hours. Cell transfection was carried out using TurboFect reagent (Thermo Fisher Scientific) according to manufacturer’s instructions. For BROX knock down, a siRNA specific to BROX (siRNA ID: 123905, CGACUCAUAGAAGCAUACGtt, Thermo Fisher Scientific) or a non-targeting control (AM461, Thermo Fisher Scientific) were used. Cells were processed for immunofluorescence 24 hours after transfection.

### Immunofluorescence and image acquisition

HEK293 cells or i^3^Ns were seeded on a poly-D-Lysine coated coverslip at the density of 300 cell/mm^2^. Cells were fixed with 4% paraformaldehyde for 15 min and, after three washes with 1x PBS, permeabilized with 0.2% Triton-X 100 for 10-15 min. Cells were blocked with 5% bovine serum albumin for 45 min and hybridized with the appropriate antibodies (see Table 1) overnight at 4 °C. Anti-mouse, anti-goat, anti-chicken, and anti-rabbit donkey secondary antibodies conjugated with either Alexa Fluor 647, Rhodamine X-red, Alexa Fluor 555, or Alexa Fluor 488 (Jackson Immunoresearch and Thermo Fisher Scientific) were hybridized for 1 h at room temperature. Coverslips were mounted onto a glass slide using Prolong Gold mounting medium (Thermo Fisher Scientific) and imaged using a widefield microscope (Leica DMi8 Thunder) equipped with a cooled CMOS camera (DFC9000 GTC). Images were acquired as Z-stacks (0.21 μm step size) using a x63 lens.

**Table 1.**
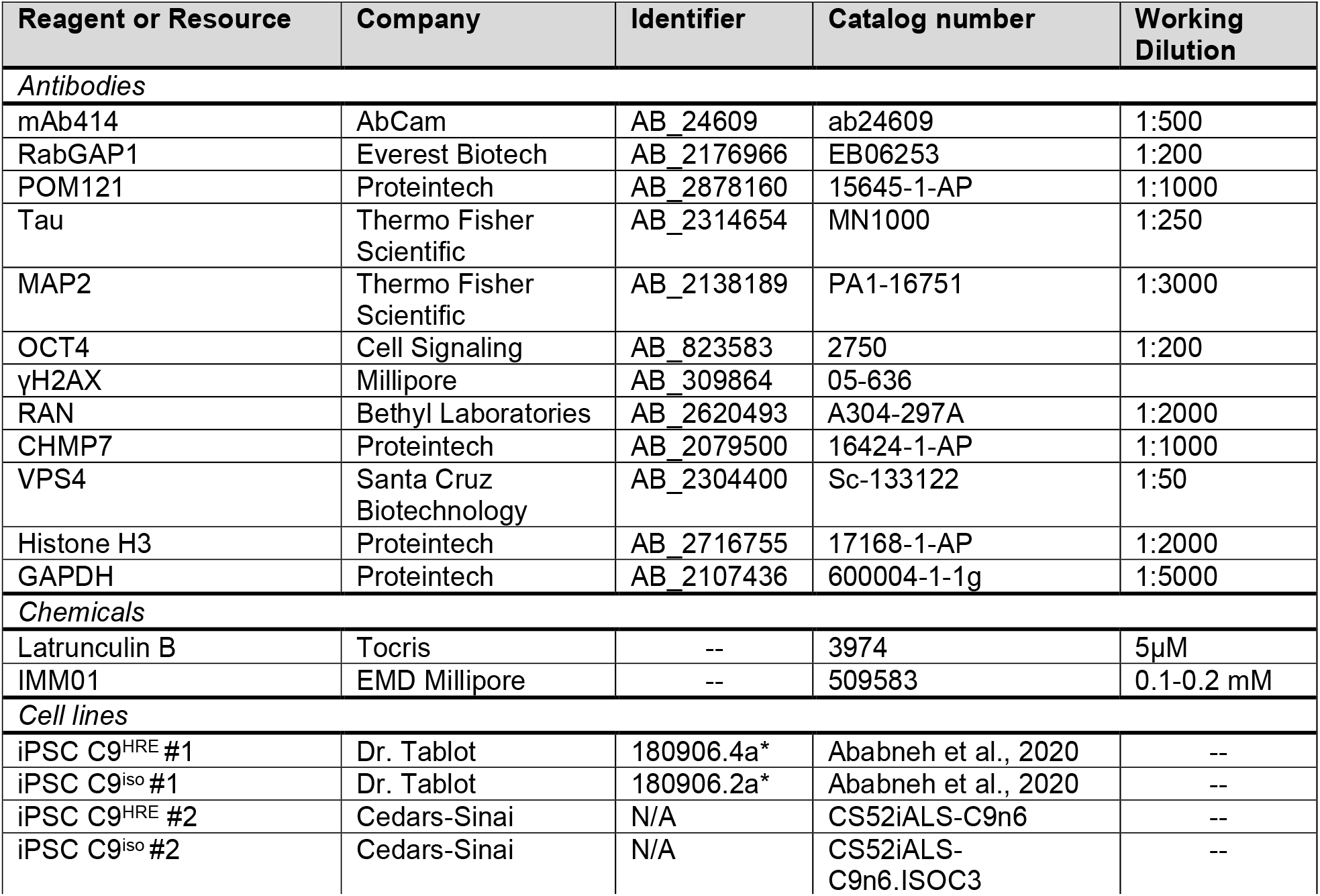
List of critical reagents or resources.

| Reagent or Resource | Company | Identifier | Catalog number | Working Dilution |
| --- | --- | --- | --- | --- |
| *Antibodies* |  |  |  |  |
| mAb414 | AbCam | AB_24609 | ab24609 | 1:500 |
| RabGAP1 | Everest Biotech | AB_2176966 | EB06253 | 1:200 |
| POM121 | Proteintech | AB_2878160 | 15645-1-AP | 1:1000 |
| Tau | Thermo Fisher Scientific | AB_2314654 | MN1000 | 1:250 |
| MAP2 | Thermo Fisher Scientific | AB_2138189 | PA1-16751 | 1:3000 |
| OCT4 | Cell Signaling | AB_823583 | 2750 | 1:200 |
| γH2AX | Millipore | AB_309864 | 05-636 | |
| RAN | Bethyl Laboratories | AB_2620493 | A304-297A | 1:2000 |
| CHMP7 | Proteintech | AB_2079500 | 16424-1-AP | 1:1000 |
| VPS4 | Santa Cruz Biotechnology | AB_2304400 | Sc-133122 | 1:50 |
| Histone H3 | Proteintech | AB_2716755 | 17168-1-AP | 1:2000 |
| GAPDH | Proteintech | AB_2107436 | 600004-1-1g | 1:5000 |
| *Chemicals* |  |  |  |  |
| Latrunculin B | Tocris | -- | 3974 | 5μM |
| IMM01 | EMD Millipore | -- | 509583 | 0.1-0.2 mM |
| *Cell lines* |  |  |  |  |
| iPSC C9<sup>HRE</sup> #1 | Dr. Tablot | 180906.4a* | Ababneh et al., 2020 | -- |
| iPSC C9<sup>iso</sup> #1 | Dr. Tablot | 180906.2a* | Ababneh et al., 2020 | -- |
| iPSC C9<sup>HRE</sup> #2 | Cedars-Sinai | N/A | CS52iALS-C9n6 | -- |
| iPSC C9<sup>iso</sup> #2 | Cedars-Sinai | N/A | CS52iALS-C9n6.ISOC3 | -- |

### Image analysis

Immunofluorescence images were deconvolved using 10 interations of an adaptative blind deconvolution algorithm (Autoquant X3, Media Cybernetics) before analysis. Fluorescence intensity levels were calculated using Image J software ^72^ according to the following protocol: stacks of our images were used to generate max intensity or sum intensity projections. A mean filter (radius 2.0 pixels) was applied to the DAPI nuclear channel to discretely separate single nuclei. Using this approach, a region of interest (ROI) was established to gate individual nuclei and measure mean fluoresce intensities (mFI) of target proteins. For quantification of nucleoporin intensity at the NE, a band 10 pixel wide was generated based on the DAPI ROI, and mFI for the indicated proteins was measured. A blind analysis was performed to qualitatively assess RanGAP1, POM121, mAb414 localization pattern on 3D stacks of individual optical slices, and to quantify the frequency of DNA leaks. To avoid misinterpretation of staining profiles, no 3D to 2D compression of the images was performed for these applications, as previously described ^26,73^.

Abnormal staining was considered if the signal was not uniformly distributed around the nucleus with the presence of empty bare segments. DAPI was used as a reference for nuclear boundary. For all experiments, raw values were normalized to the mean of the control condition.

For FRET assays, HEK293 cells were transfected with the tension sensor (TS) ^74^, mVFP ^75^, or mTFP ^76^ plasmids 24 hours before imaging. Cells were then treated for 5 hours with the formin agonist IMM01 (100 or 200 μM) or for 24 hours with the actin depolymerizing drugs Latrunculin B (5 μM) to either increase or decrease actin polymerization. For live cell imaging, cells were treated with IMM01 and imaged immediately after to monitor changes to FRET efficiency over the course of 1 hour. Cells were then fixed and processed for fluorescence imaging as previously described ^51,74^. FRET efficiency was quantified using the ImageJ PixFRET plug-in using the Acceptor Model ^77^. Higher FRET values are indicative of decreased mechanical tension or increased compressive force on the nucleus.

### Protein fractionation and Western Blot assays

For whole protein extraction, cells were centrifuged at 250 *x* g for 10 min at 4°C and rinsed once with 1x PBS. Pellets were lysed with lysis buffer (10 mM Tris-HCL pH 8, 100 mM NaCl, 1%, 1 mM EDTA pH 8, 1%NP40) supplemented with protease inhibitor (Complete EDTA-free, Roche) and phosphatase inhibitor (phosphoSTOP, Roche). Samples were sonicated at the frequency of 40 kHz continuously of 10s (Sonifier® SFX150, Emerson). For nucleocytoplasmic fractionation, cells were lysed for 30 min in a sucrose buffer (0.32 M sucrose, 0.5% NP40, 3 mM CaCl2, 2 mM magnesium acetate, 10 mM Tris-HCL pH 8, 1 mM dithiothreitol, 0.1 mM EDTA) supplemented with protease inhibitor and centrifuged at 1500 x *g* for 5 mins at 4°C. The supernatant was removed and centrifuged at 2600 x *g* for 5 mins at 4°C and saved as the cytoplasmic fraction. The pellet containing the cells’ nuclei was washed in sucrose buffer and centrifuged at 2600 x *g* for 5 mins at 4°C. The pellet was then resuspended in a lysis buffer (20 mM Tris, 150 mM NaCl, 1% Triton X100) supplemented with protease inhibitor, and sonicated at the frequency of 40 kHz continuously for two rounds of 10s. Protein concentrations were measured by Bradford protein assay. For each sample, 30 μg of protein extract was diluted in Laemmli buffer (60 mM Tris-Cl pH 6.8, 2% SDS, 10% glycerol, 5% beta-mercaptoethanol, 0.01% bromophenol blue) and then resolved by SDS-PAGE on Mini 4% - 20% Novex Tris-glycine gels (Thermo Fisher Scientific) and transferred onto nitrocellulose membranes (Thermo Fisher Scientific). Membranes were blocked with Odyssey Blocking Buffer (LI-COR) and probed with primary antibodies overnight at 4°C (see Table 1). Secondary antibodies conjugated with IRDye® infrared fluorophores (LI-COR) were incubated for 1 h at room temperature. Blots were visualized using the Odyssey Infrared Imaging System (LI-COR).

### Statistical analysis

Statistical analyses were performed using Prism 9 software package (GraphPad). Normality of the samples was assessed using the D’Agostino & Pearson Omnibus test. According to normality, parametric or non-parametric tests were used to assess significance, defined as p < 0.05.

## ACKNOWLEDGEMENTS

This work was supported by the National Institute of Neurodegeneration and Stroke (NINDS) within the National Institute of Health (NIH) under Grant # R01NS116143 to CF and the RI-INBRE Early Career Development Award to CF. SS was the recipient of a fellowship from the PhD program in Experimental Medicine, Università degli Studi di Milano. AR acknowledges “Aldo Ravelli Center for Neurotechnology and Experimental Brain Therapeutics”, Università degli Studi di Milano and is supported by a grant from the Italian Ministry of Health (RF-2019-12368778). Access to core facility was made through the Institutional Development Award (IDeA) Network for Biomedical Research Excellence from the National Institute of General Medical Sciences (NIGMS) of the NIH under grant # P20GM103430. We are indebted to Dr. Ward (NIH) and Dr. Talbot (University of Oxford, UK) for sharing the *C9ORF72* mutant and isogenic control iPSC lines, and to Dr. Agromayor for sharing the BROX WT and L350A plasmids. The pcDNA Nesprin TS and mCherry-DN KASH plasmids were a gift from Daniel Conway (Addgene #68127 and #125553). mVenus C1 (Addgene #27794) was a gift from Robert Campbell and Michael Davidson. mTFP1-N1 (Addgene #54521) was a gift from Steven Vogel. CLYBL-TO-hNGN2-BSD-mApple was a gift from Michael Ward (Addgene plasmid # 124229).

