## Supplementary Material for "Altered nuclear envelope homeostasis is a key pathogenic event in C9ORF72-linked ALS/FTD"

### Content:

- Figure S1. *$i^3$PSC differentiation and maturation to cortical-like neurons.*
- Figure S2. *Time-dependent reduction in nucleoporin levels at the NE of $i^3$Ns.*
- Figure S3. *FRET analysis of nuclear tension in HEK293 cells by live cell imaging.*
- Figure S4. *$BROX^{L350A}$ does not rescue NPC injury.*
- Table S1. *iPSC lines used in this study.*
- Reference List for Supplementary material

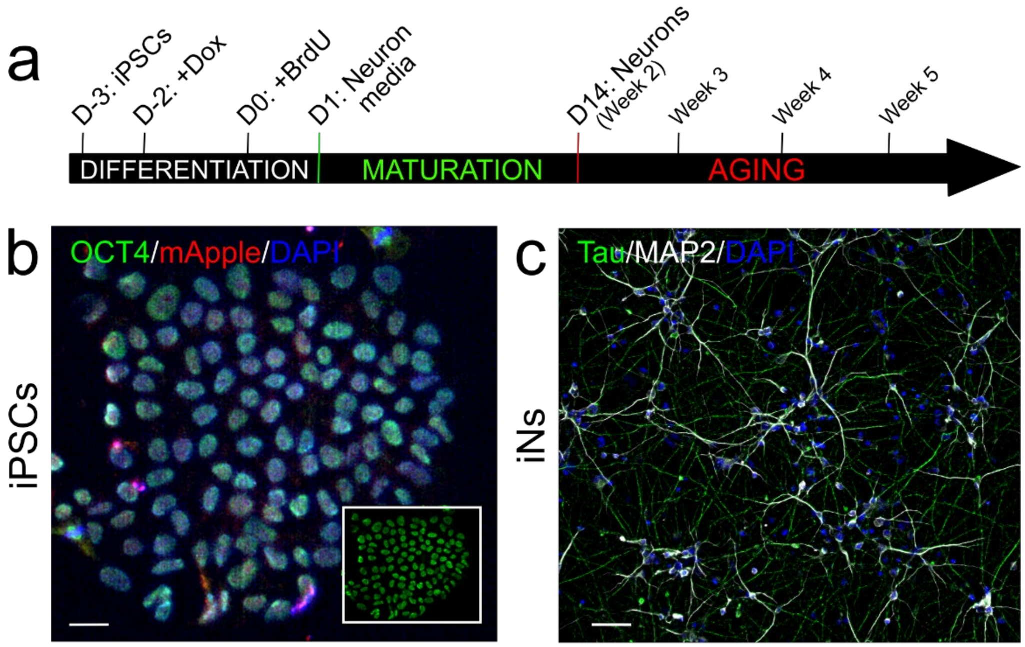


**Figure S1. i$^3$PSC differentiation and maturation to cortical-like neurons.** *Related to Figure 1*. **a**. Schematics of iPSC differentiation to cortical-like i$^3$ neurons (i$^3$Ns). i$^3$PSCs are treated with doxycycline to induce the expression of the transcription factor NGN2. Two days after induction, neuroprogenitors are plated on poly-lysine coated coverslips in neuron media and matured for a minimum of 2 weeks. **b-c**. Representative images of i$^3$PSCs (*b*) and i$^3$Ns (*c*) labeled with stage-appropriate markers. OCT4 (*green, b*) labels stem cells, while Tau (*green*, *c*) and MAP2 (*greys*, *c*) are neuronal-specific makers. DAPI (*blue*) was used to labels the cells' nuclei. mApple (*red* in *b*) is a marker of the differentiation cassette [1]. Scale bar: 25μm in *b*, 50μm in *c*.

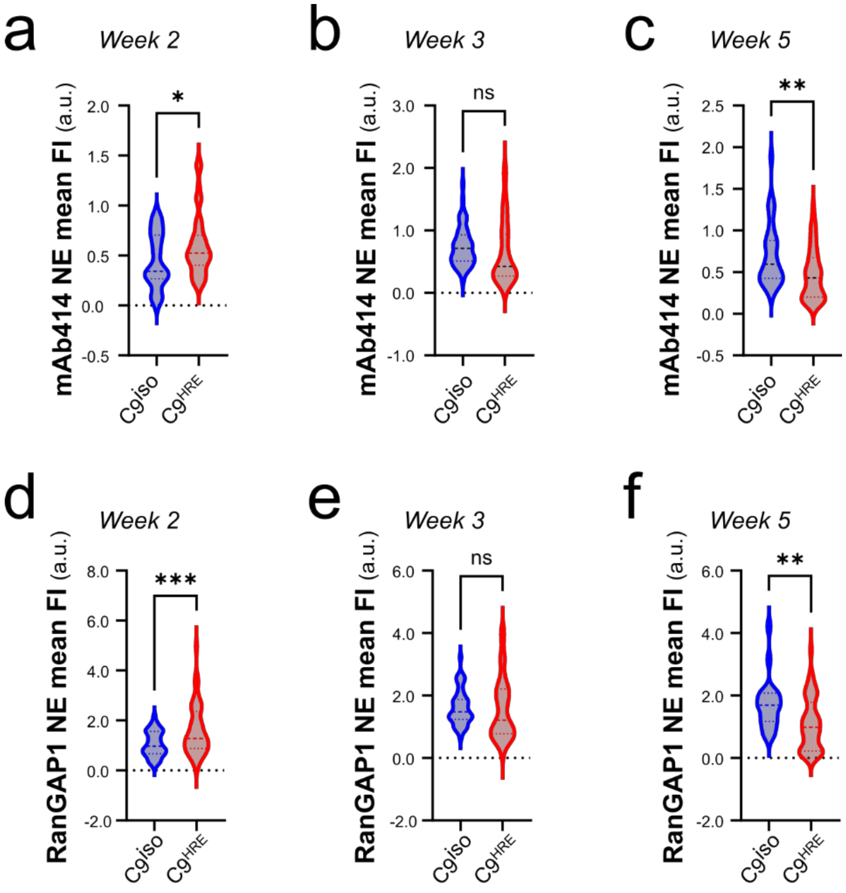


**Figure S2. Time-dependent reduction in nucleoporin levels at the NE of i$^3$Ns.** *Related to Fig. 1.* Quantification of NPC markers mAb414 (***a-c***) and RanGAP1 (***d-e***) at the NE in i$^3$Ns at 2, 3, and 5 weeks *in vitro*. Mutant C9$^{HRE\#1}$ neurons show slightly elevated levels of both markers at 2 weeks of age compared to C9$^{iso\#1}$ controls. This change is followed by a gradual decrease in NE levels of both markers, becoming statistically significant at week 5 (violin plots show data distribution; dashed lines are mean and SD; Welch's *t* test, n=22 and 47 for C9$^{iso\#1}$ and C9$^{HRE\#1}$, respectively, * $p$<0.05; ** $p$<0.01; *** $p$<0.001).

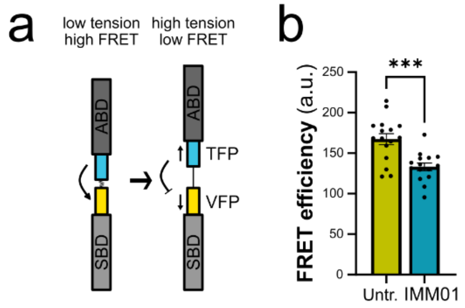


**Figure S3. FRET analysis of nuclear tension in HEK293 cells by live cell imaging. a.** Schematic representation of the FRET sensor used[2,3], based on the LINC complex protein Nesprin2. When expressed in cells, the sensor is incorporated in existing LINC complexes by binding to SUN proteins via the SUN binding domain (SBD). The actin-binding domain (ABD) allows the sensor to directly bind actin. A FRET pair of fluorescent proteins (i.e., VFP and TFP) is located in between the two domains and is separated by a molecular spring. Cytoskeletal tension will pull the two fluorescent proteins apart, reducing FRET efficiency. Vice versa, low cytoskeletal tension will allow for increased FRET due to the proximity of the FRET pair. **b.** Quantification of FRET efficiency in HEK293 cells treated with 0.1mM IMM01 and immediately imaged by live cell imaging. Bars are mean and SEM (Student's t test; n=15 and 19 for Untr. and IMM01, respectively; *** $p$<0.001).

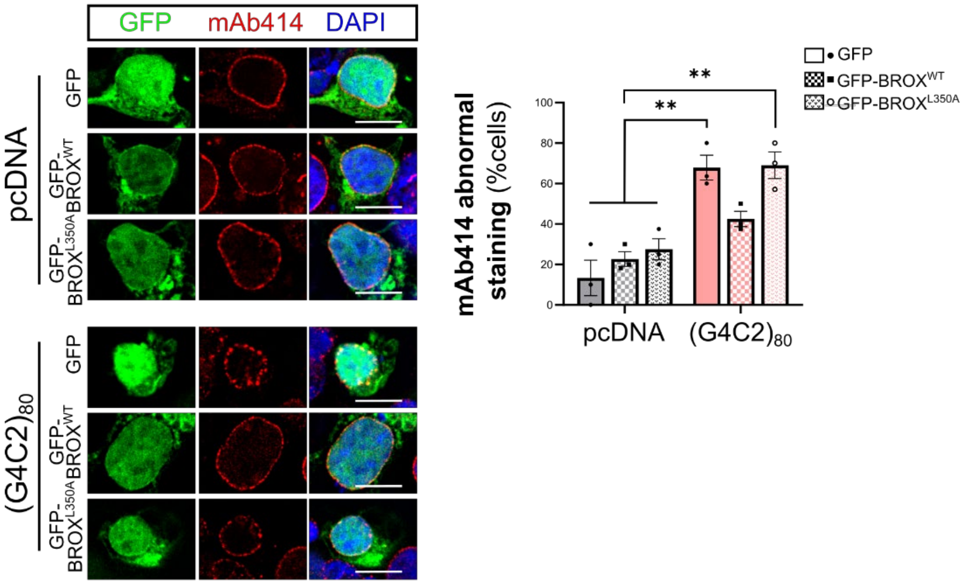


**Figure S4. BROX$^{L350A}$ does not rescue NPC injury.** *Related to Figure 5.* Representative images and quantification of NPC injury (mAb414, *red*) in $(G4C2)_{80}$ or control (pcDNA) HEK293 cells expressing GFP-tagged wild type BROX (GFP-BROX$^{WT}$), mutant BROX (GFP-BROX$^{L350A}$), or GFP alone (*green*). While GFP-BROX$^{WT}$ leads to significant rescue of NPC disruption, the effect is abrogated by the L350A mutation, which impairs its binding to Nesprin2 [4]. Bars represent mean and SEM; two-way ANOVA; n=3; ** $p$<0.01. Scale bars: 10µm.

| Line # / ID in this study | Parent Tissue | Mutation | AS | AO | Gender | Reference/ Source |
|---|---|---|---|---|---|---|
| 180906.4a / $C9^{HRE\#1}$ | Fibroblast | 6 kb HRE | 62 | N/A | Female | [5,6] |
| 180906.2a / $C9^{iso\#1}$ | -- | Isogenic ctrl of 180906.4a | -- | N/A | Female | [6] |
| CS52iALS-C9n6 / $C9^{HRE\#2}$ | Fibroblast | 6-8 kb HRE | 49 | 57 | Male | Cedar Sinai |
| CS52iALS-C9n6.ISO / $C9^{iso\#2}$ | -- | Isogenic ctrl of CS52iALS-C9n6 | -- | -- | Male | Cedar Sinai |

**Table S1. iPSC lines used in this study.** AS: age at sampling; AO: age of onset; HRE: hexanucleotide repeat expansion
